# Cost-benefit analysis of cryogenic electron tomography subtomogram averaging of chaperonin MmCpn at near atomic resolution

**DOI:** 10.1101/2024.01.30.577584

**Authors:** Yanyan Zhao, Michael F. Schmid, Wah Chiu

## Abstract

Cryogenic electron microscopy single particle reconstruction (cryoEM-SPR) has evolved into a routine approach for determining macromolecule structures to near-atomic resolution. Cryogenic electron tomography subtomogram averaging (cryoET-STA) towards similar resolution, in contrast, is still under active development. CryoET can capture the 3D snapshot of individual macromolecules by stage tilting, offering multiple angular views per particle than the single particle reconstruction approach. Here we use the archaea chaperonin MmCpn as a model macromolecule to investigate the resolution limiting factors of cryoET-STA in terms of cumulative electron dose, ice thickness, subtomogram numbers and tilt angle ranges. By quantitative analysis of these factors against the STA reconstruction resolution, we delineate the feasibility of attaining high resolution structure determination with cryoET-STA. This study provides biophysical guidance for the application of cryoET-STA towards high resolution and the cost against benefit of using cryoET-STA to achieve an efficient outcome at the desired resolution.

## Introduction

CryoEM-SPR allows the structural study of purified and concentrated macromolecules in solution at near to full atomic resolution. In cryoEM-SPR, the specimen embedded in a thin layer of vitreous ice is exposed to a cumulative electron dose throughout the acquisition process, producing a single 2D projection image per acquisition area. Alignment, classification, and reconstruction of a sufficient number of 2D projection images (tens of thousands to millions) of randomly oriented macromolecules leads to 3D density maps. With the advances in electron microscope hardware, cameras, and image-processing algorithms, the cryoEM-SPR approach has become a useful and routine method to investigate the structures and conformational dynamics of a broad range of biochemically purified biological macromolecules at near to full atomic resolution ^1,2^.

In contrast, cryoET-STA is an imaging method often used to visualize macromolecules and subcellular structures within their cellular environment. In cryoET, the specimen is physically rotated around a tilt axis during data acquisition and a tilt-series of 2D projections of the exposed area is collected from multiple angles. This tilt-series is then computationally aligned and reconstructed to yield a tomogram (i.e., a 3D density map) of the object being imaged. The abundance of the target macromolecule *in situ*, however, is usually orders of magnitudes lower than the concentrated specimen after purification. Hence only hundreds to thousands of copies of 3D subtomograms of target macromolecules in the tomograms are generally extracted, computationally aligned and averaged to a subnanometer resolution 3D density map. In some recent cases, a large dataset of tomograms with highly abundant macromolecules either *in situ* or purified can also yield a near-atomic resolution density map by cryoET-STA ^3,4,5,6^.

While cryoEM-SPR is still under active development particularly on software to sort out heterogeneous particle images, cryoET-STA has been conceived as a powerful technique to reveal the molecular sociology within the cellular environment ^7^. The trend of pursuing subnanometer resolution structural determination by cryoET-STA is growing rapidly ^8,9,10,11^. However, the cost-effectiveness of the cryoET-STA over cryoEM-SPR to resolve the macromolecules becomes a commonly asked question, given the time and computational complexity associated with cryoET-STA. In this study, we examine several resolution-limiting factors in cryoET-STA, including ice thickness, macromolecule number, electron damage, and tilt angle range. With this benchmark study, we provide a useful insight into the costs and benefits in current applications of cryoET-STA either in purified or *in situ* macromolecules.

## Results

### Reconstruction of MmCpn in closed conformation by cryoET-STA

Apoferritin, ribosome or virus have been reported with high resolution cryoET-STA ^4,5,6^. Structural rigidity, high symmetry, or large molecular weight (high signal to noise ratio) contributes collectively to the subtomogram alignment accuracy of these macromolecules. To understand how subtomogram averaging applies to protein complexes other than these ideal targets, we chose to investigate cryoET-STA methodology using the archaea chaperonin MmCpn. MmCpn is known as a hexadecameric homo-oligomeric protein complex with molecular weight of one megadalton ^12^. The complex adopts a closed conformation with D8 symmetry under the ATP and AlFx condition. In this study, we collect around 500 tilt series with a total of 170,000 apparent MmCpn macromolecules on an EM grid from -18 to 18 degree tilt range (Figure 1a). The data is processed following the workflow (Figure 1b), yielding a 2.56 Å STA map of MmCpn in closed conformation by imposing D8 symmetry, from a total of 158,666 hexadecameric MmCpn macromolecules (Figure 1c and Figure S1a). Surprisingly, we also observe a small population of MmCpn (7% of the total macromolecules) in octadecameric state with D9 symmetry that has not previously been reported (Figure S1b). The STA reconstruction resolution is estimated by the FSC gold standard with two split sets of subtomograms. In each subunit of the D8 symmetry reconstruction, an ADP coordinated by AlF_3_ and Mg^2+^ was observed in the nucleotide binding pocket as well as a water molecule at the expected position. Furthermore, an extra density at the opposite side of the ADP is detected, possibly belonging to the K^+^ ion as shown in previous X-ray crystallographic study of group I chaperonin GroEL^13^ (Figure 1e). We further characterize the resolvability of the local residues by Q score ^14^ as expected for the claimed resolution (Figure 1d).

**Figure 1.**
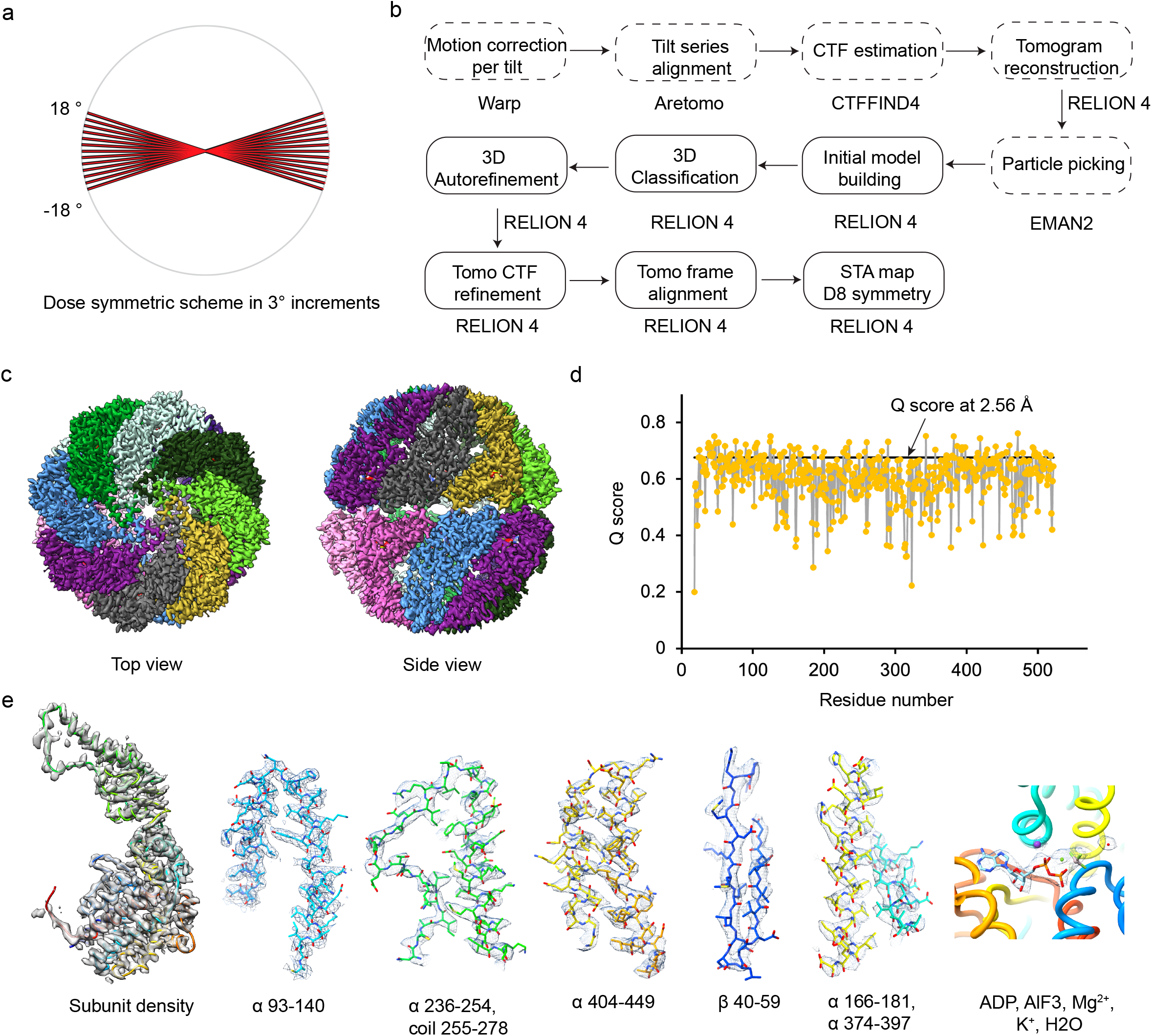
(a) The tilt series is collected in a dose-symmetric manner to cover from -18° to 18° in 3° increments. (b) The data processing workflow with various software packages. The input to the data processing steps shown in the dashed boxes are individual tilt series or reconstructed tomograms, while input to the rest of the steps are individual subtomograms from multiple tomograms. (c) Reconstruction of hexadecameric MmCpn in the closed conformation with D8 symmetry. (d) Q score plot of the local amino acid residue resolvability in the single subunit of the reconstruction shown in (c). (e) Segmented MmCpn subunit density (left) from a 2.56 Å reconstruction with D8 symmetry and the local side chain resolvability at various secondary structure regions and the ligand bound in thenucleotide binding pocket (right). The K^+^ ion and Mg^2+^ ions are colored in purple and green while the water is colored in red.

### Radiation damage analysis of different cumulative electron doses throughout the tilt series collection

The dose accumulation and increased ice thickness both contribute to the impaired resolution at high degree tilts generally recorded in the later part of a tilt series collection. In our data, 94% of hexadecameric MmCpn macromolecules are embedded in vitreous ice thinner than 1200 Å (see later figure). The ice thickness from 0 to 18 degrees only increases by up to around 60 Å. Therefore, ice thickness varies little within a tilt series collection at these angles.

To investigate the major resolution loss of the data in all the tilt series due to the accumulated dose damage, we reconstruct the MmCpn map with D8 symmetry using the same set of 158,666 macromolecules except only at one tilt angle at a time (Figure 2a, Figure S2). By plotting the map resolution against the cumulative dose at any specific tilt, we observe a linear loss of high resolution structural information as the cumulative dose increases (Figure 2b). The 2 to 3 Å signal resolving small side chains is quickly lost upon the exposure of 6 electrons/Å^2^ while the 3 to 4 Å signal resolving bulky side chains is lost at 12 electrons/Å^2^. As the dose further accumulates in the course of a tilt series, we observe a significant damage to the secondary structural elements (alpha helices and beta strands) ^15^. First, the beta strand densities are no longer resolved due to the loss of 4-5 Å signals before the cumulative dose reaches 20 electrons/Å^2^. Next, the alpha helices (resolvable at 8 Å) are further damaged as the sample is exposed to a total dose of 30 electrons/Å^2^. This observation demonstrates electron dose significantly and systematically damages the side chains and then secondary structural elements of a protein molecule within the course of total exposure of 30 electrons/Å^2^ (Figure 2c, Table 1). Given the minimal increase in the ice thickness from +/-15 degree stage tilting, the dose symmetric data collection scheme we used in this study has no advantage over a continuous scheme in acquiring the high resolution tilts within 30 electrons/Å^2^.

**Table 1.** The averaged Q score of local side chain (sc), backbone (bb) and amino acid residue of the subunit for each reconstruction at the specific cumulative dose and estimated resolution. The expected Q score is obtained from all of the PDB structures reported at that resolution.

|  | 3 e/Å<sup>2</sup> | 12 e/Å<sup>2</sup> | 21 e/Å<sup>2</sup> | 30 e/Å<sup>2</sup> | 39 e/Å<sup>2</sup> |
| --- | --- | --- | --- | --- | --- |
| Resolution (Å) | 2.56 | 3.82 | 6.57 | 7.96 | 10.42 |
| Q score (sc) | 0.58 | 0.27 | 0.10 | 0.05 | 0.02 |
| Q score (bb) | 0.62 | 0.44 | 0.21 | 0.07 | 0.07 |
| Q score (Residue) | 0.603 | 0.355 | 0.154 | 0.06 | 0.051 |
| Q score (Residue) expected | 0.68 | 0.44 | 0.19 | 0.15 | 0.10 |
| Structure features damage | small sc | bulky sc | beta strands | alpha helices | all sc and secondary structures |

**Figure 2.**
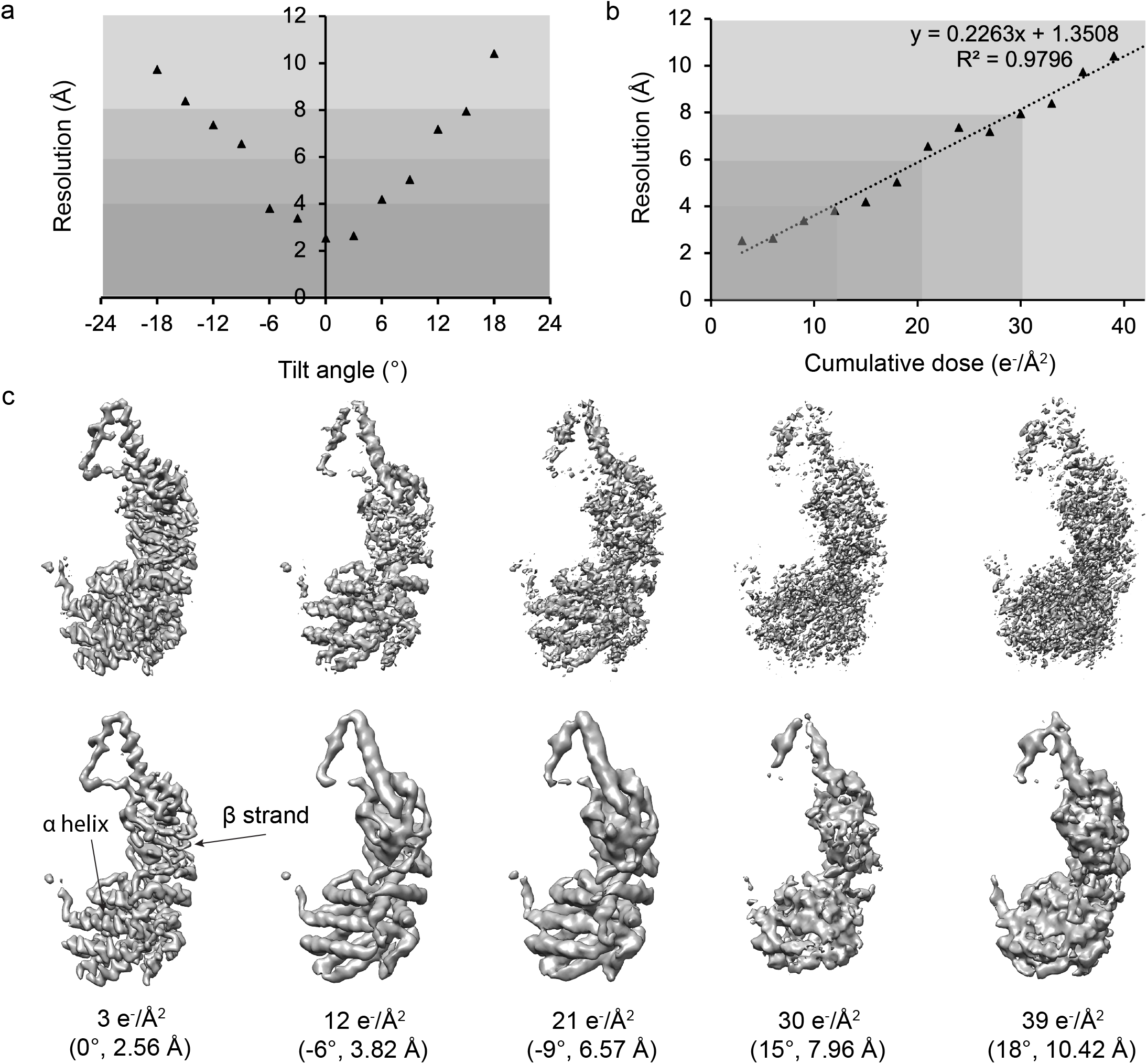
(a) The resolution of reconstructions from each specific tilt angle with the same set of 158,666 macromolecules is plotted against the tilt angle. (b) The resolution of the reconstruction from each tilt in (a) is plotted against the cumulative dose (electron/Å^2^) in the tilt. (c) Top row shows the segmented MmCpn subunit density from the maps reconstructed from the subtomograms at a specific tilt angle. Bottom row shows the gaussian filtered maps from the top row.

### Number of tilts usable for subtomogram averaging at different target resolutions

In cryoEM-SPR, each particle image contributes only one central slice to the 3D Fourier space. In contrast, in subtomogram averaging, each subtomogram containing a single macromolecule in a 3D tilt series contributes a series of central slices in 3D Fourier space, providing more information from different angles per subtomogram than a single image in cryoEM-SPR. Thus, one may raise the question whether subtomogram averaging can yield a similar resolution map with fewer macromolecules than the single particle reconstruction approach. However, due to the electron damage, the individual tilt images within a tilt series contribute structural information unequally. Because of the linear resolution loss shown in the above study as the electron dose accumulates during a tilt series collection (Figure 2b), the effective number of tilts contributing constructively is linearly and inversely related to the better resolution (Figure 3).

**Figure 3.**
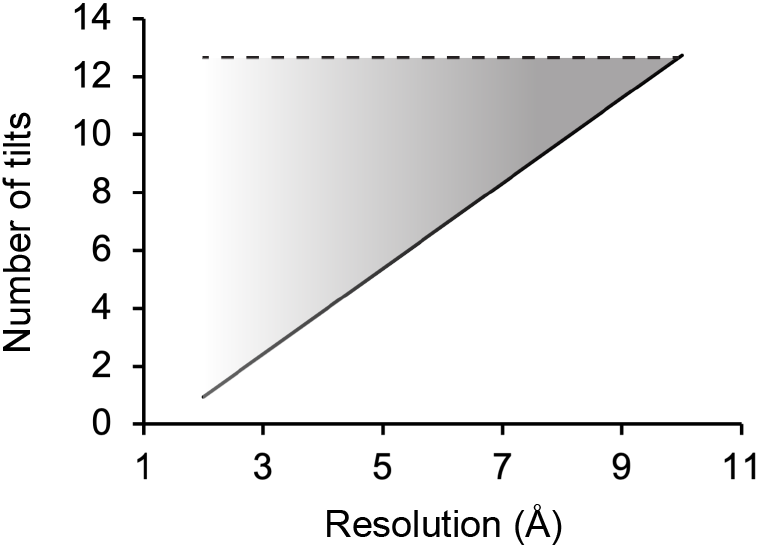
The dark line indicates the number of tilt images within our tilt series that contributed constructively to the target resolution as a result of dose constraints. The shaded area signifies the potential improvement that could be made to increase the number of usable tilts for the target resolution. The dashed line represents the ideal scenario if there was no electron damage.

For example, if the resolution is aimed for 8 Å, the first 10 tilts would contribute structural information at 8 Å and thus can be considered to be useful tilts towards this target resolution. In contrast, if the target resolution is 2.6 Å, only 2 tilts contribute effectively to that resolution because the rest of the tilts contain only lower resolution content due to radiation damage (Figure 3). Therefore, a medium resolution target (e.g., 8 Å) would benefit from the multiple tilts by subtomogram averaging with fewer macromolecules used than in cryoEM-SPR. On the other hand, a near-atomic resolution target (e.g., 2.6 Å) by subtomogram averaging would require half the number of macromolecules used in single particle reconstruction (i.e. two usable tilts per macromolecule). However, if more of the tilt images in a tilt series could offer high resolution data, we could further reduce the number of macromolecules needed towards this resolution target. This could be possibly achieved in the future by less electron damage with lower temperature cooling ^15,16,17^, higher contrast with phase plate^18^, better DQE with new detectors^19^, or accurate alignment of noisy images under lower dose by yet to be developed image processing algorithms.

### Effect of the ice thickness in the subtomogram averaging

In recent years, the *in situ* reconstructions of macromolecules from different types of cells by subtomogram averaging are reported ^3,4,5,8,9,10,11^. The ice thickness of imaged cellular samples varies a lot depending on acquisition area and method of data acquisition and the type of cell, with an ice thickness at ∼1500 Å from focus ion beam milled samples. However, how the different ice thickness of the sample affects the resolution in cryoET-STA remains unclear. Here we plot the macromolecule distribution and measure the ice thickness of all of our ∼500 tomograms in 3D (see Methods), and observe a large variation of ice thickness within and across the tomograms (Figure 4a, Figure S3). In many of our tomograms, MmCpn embedded in the vitreous ice overlapping in the electron path and considered to be useless by cryoEM-SPR, are clearly separable in 3D space and usable by cryoET-STA.

**Figure 4.**
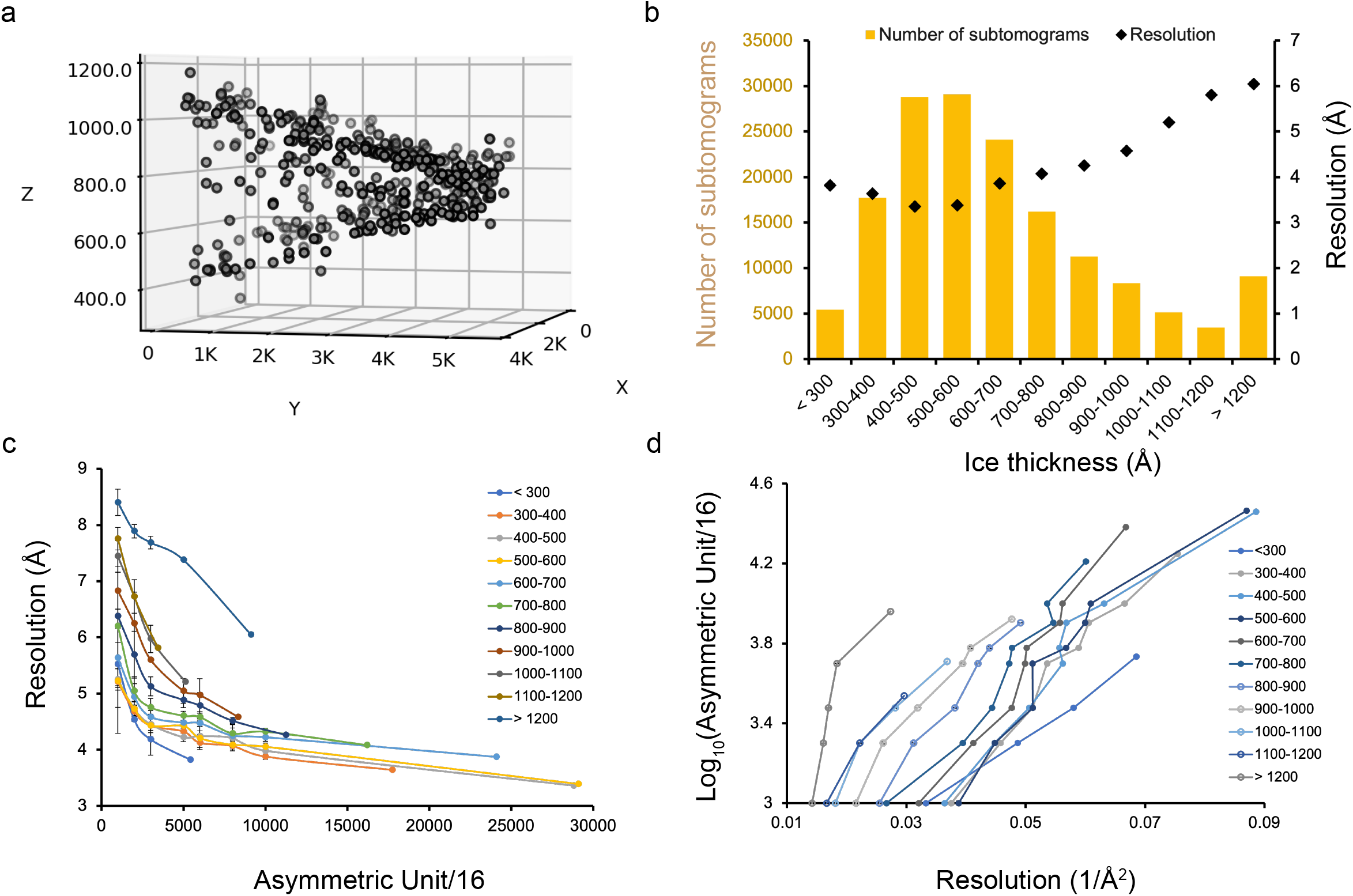
(a) An example of the uneven macromolecule distribution in a tomogram demonstrates the non-uniform ice thickness in the acquired area (X,Y,Z axis in pixel with sampling at 1.08 Å/pixel). (b) The histogram shows the statistics of the ice thickness per MmCpn (left Y axis) from all the tomograms in the dataset. The black points represent the reconstruction resolution (right Y axis) from the corresponding subtomogram subsets in each ice thickness bin. (c) The resolution of reconstructions from subsampled subsets of each ice thickness bin is plotted against the subsampling size. The x axis denotes subtomogram number for the hexadecameric chaperonin. (d) From (c), the log_10_ of the subsampling size is plotted against the inverse of the squared resolution.

Previous studies roughly estimate the sample ice thickness and assign a single ice thickness value to a single image acquisition ^20,21^. To gain a more accurate insight into how the ice thickness across the imaging area affects the resolution in the STA maps, we measure the local ice thickness of an acquisition area and assign the nonuniform ice thickness values to individual macromolecules of the tomogram. Our analysis in our dataset shows that MmCpn complexes are embedded in the ice with thickness ranging from less than 300 Å to over 1200 Å (Figure 4b). We reconstruct an individual 3D map from each ice thickness class of subtomograms and evaluate the relationship of subtomogram number to the resolution for different ice thicknesses. We find that similar numbers of subtomograms from different ice thickness classes yield reconstructions at significantly different resolutions.

For example, around 5,000 macromolecules embedded in a thin (< 300 Å) ice layer are reconstructed to 3.8 Å while a similar number of macromolecules from 1000 to 1100 Å thick ice yield a 5.2 Å reconstruction. To accurately assess the correlation between the number of subtomograms and the resolution for each ice thickness, subtomograms from each ice thickness class are subsampled to multiple subsets of various sizes and reconstructed to estimate the resolution (Figure 4c). This analysis shows that the increase of the subtomogram number generally improves the resolution more significantly in the low to medium resolution range (10 to 4 Å) than at near-atomic resolution (4 to 2 Å) for the ice thickness ranging from 300 to more than 1200 Å (Figure 4c, 4d).

The decrease of elastic scattering in thick ice dampens the envelope function quickly and more subtomograms are needed to reach high resolution. We therefore examine the reconstructed maps with fewer than 6,000 macromolecules from different ice thickness classes (Figure 5). If the ice thickness is less than 300 Å, we only need 3000 to 5414 macromolecules to achieve 4.2 Å and 3.8 Å resolution maps, respectively (Figure 5a, Figure 4c). At these resolutions, beta strands can be confidently modeled. For ice thickness between 300 Å and 800 Å, a similar number of subtomograms allows reconstructions better than 5 Å, with beta sheets reasonably detectable (Figure 5b, Figure 4c). Even the samples with 1000 Å thick ice still allow structure determination better than 6 Å resolution readily (Figure 4c). In contrast, ice thicker than 1200 Å significantly impairs the medium to high resolution structure determination; nevertheless, 6 to 7 Å reconstruction is still achievable with a reasonable number of subtomograms (Figure 5c, Figure 4c). These observations explain many recent reports of low to medium resolution structures (10 to 6 Å) by cryoET-STA to reconstruct the macromolecules embedded in ice of 1500 - 2000 Å from FIB milled lamellas^10,22^.

**Figure 5.**
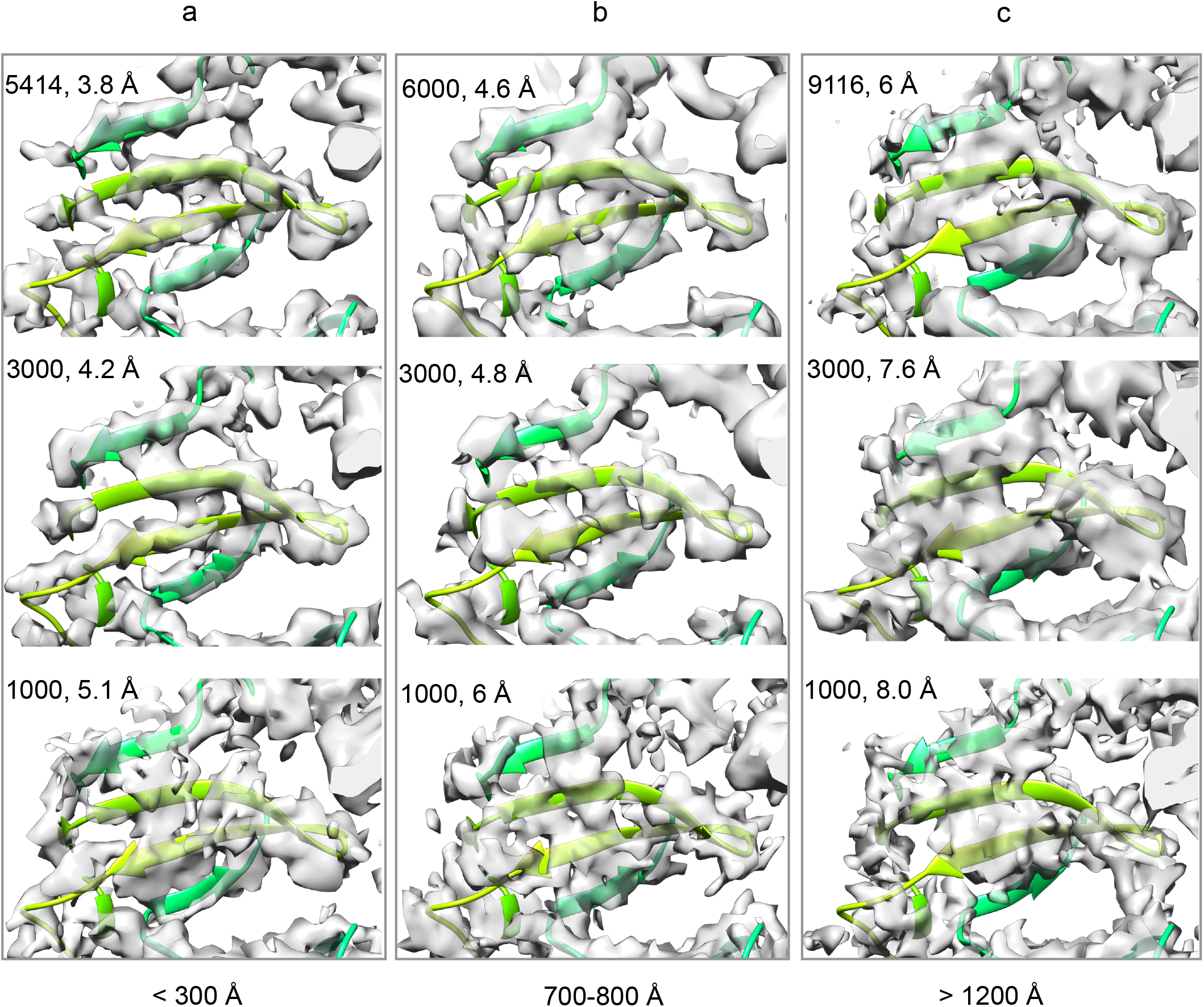
The resolvability at the beta-sheet region of MmCpn reconstruction from three different numbers of subtomograms yielding various resolutions (both indicated at the top left corner of each panel) for three ice thickness bins. Column (a) represents the reconstructions from macromolecules embedded in ice thinner than 300 Å, column (b) for ice thickness between 700 and 800 Å, and column (c) for ice thickness over 1200 Å.

## Discussion

In this study, we use the archaea chaperonin MmCpn as a model macromolecule to investigate the factors limiting the achievable resolutions in cryogenic electron tomography and subtomogram averaging. We reconstructed the Mmcpn with D8 symmetry by subtomogram averaging to 2.56 Å, close to the Nyquist limit. We investigated two major causes of resolution loss among the tilts, the electron radiation damage and the embedding ice thickness. The first 12 electrons/Å^2^ damages the near-atomic resolution information (4 Å), and a total cumulative dose of 30 electrons/Å^2^ further damages secondary structural elements (4 to 8 Å). A cumulative dose of more than 40 electrons/Å^2^ will destroy the secondary structural elements (Figure 2 and 5). We assessed the effect of embedding ice thickness on final STA resolution, and found that more macromolecules improve the resolution more efficiently at low to medium resolution range (10 to 4 Å) than at near-atomic resolution (4 to 2 Å), when the ice is thicker than 300 Å (Figure 4c, 4d).

In cryoEM-SPR, the number of particle images (expressed in log_10_ N) is linearly related to the square of spatial frequency (reciprocal of resolution square 1/Å^2^) under 10 Å with a uniform slope referred to as Rosenthal-Henderson B factor ^23^. In cryoET-STA, however, the correlation between the number of subtomograms and resolution would be nonlinear across the resolution range. The individual tilt of a tilt series contributes unequally with a gradual loss of structural information due to the electron damage across the tilts. In ice thicker than 300 Å for our benchmark macromolecule with a dimension of 148 Å by 154 Å, the slope of such plot is expected to be small at low resolution (10 to 8 Å) but gradually increase and approximate the Rosenthal-Henderson B factor by single particle reconstruction at near-atomic resolution (4 to 2 Å). The higher the resolution we seek (e.g. better than 3Å), the less advantage most of the images recorded in a later part of a tilt series offers.

While radiation damage occurs through a tilt series, cryoET-STA allows separation of individual molecules into 3D subtomograms and rescue overlapping macromolecules on the electron path. It is especially important in the cellular context where macromolecules of interest are often surrounded by other types of macromolecules or subcellular components. This study indicates another niche of cryoET-STA over cryoEM-SPR at 4 to 8 Å resolution. For example, abundance of purified macromolecules may be limited due to difficulty in biochemistry preparation or with poor distribution of macromolecules on the EM grid, or the projects require millions of macromolecules to sort out the large conformational and compositional heterogeneity. Under these circumstances, a distinct cryoEM map could be reconstructed with 4 to 10 times fewer macromolecules by STA compared to SPR. Even when the target resolution is close to 4 Å, if good distributions of macromolecules are embedded in relatively thin ice as shown here, we could expect a 75% reduction of the macromolecule number with 4 tilts in a tilt series within 12 electrons/Å^2^ and 83% reduction with 6 tilts in a tilt series within same amount of dose while maintaining the 4 Å structural information. However, the ice thickness is an additional major factor limiting high resolution determination, in particular for the FIB milled samples beyond 1500 Å. Although resolution beyond 6 Å is achievable with a reasonable to exceptionally large number of macromolecules at this ice thickness, we expect it can be further improved in the future.

Current practice in either cryoET-STA or cryoEM-SPR is limited at around one electron/Å^2^ per frame to allow accurate motion correction and tilt alignment^24^. This dose usage determines the fast resolution loss among the tilts and limits the subtomogram averaging to low to medium resolution (10 to 4 Å) for the sake of cost-effectiveness. If a lower dose per frame (or tilt) is allowed for accurate motion correction (or tilt alignment) so that more tilts within a tilt series can contribute effectively at high resolution, subtomogram averaging would advance significantly for *in situ* structural determination and become a competitive technique against the single particle reconstruction at near-atomic to full resolution. This could possibly be achieved by new algorithms offering accurate alignment at lower dose, new detector with a better DQE, or phase plate providing high image contrast. In addition, liquid helium cooling (down to 4K) reducing the radiation damage per tilt would also outweigh the benefit over cost further than the current practice^17^.

As the ice thickness varies little among the low degree tilts (−18° to 18°), the favored dose symmetric tilt series collection in current practice of cryoET-STA offers marginal advantage over other data collection schemes such as the continuous or bidirectional collection schemes. Therefore, we suggest that users may flexibly customize the tilt series collection strategy ^25,26^. Broadly speaking, this study supports the dose fractionation theory of Glaeser ^27^, with the caveat that the electron damage in a tilt series limits its resolution and practical applicability in the real world. Weighing the cost/benefit ratio against the desired resolution will help achieve better and more efficient throughput.

## Methods

### Sample preparation and freezing

Archaea MmCpn was purified as described previously^28^. The complex was incubated with ATP, Al(NO_3_)_3_, and NaF to induce the closed state at room temperature for 1hr. 3 ul sample of 1 μM MmCpn was applied on the Quantifoil R2/1 EM grid and flash frozen in liquid ethane using Vitrobot.

### Data collection

Sample was imaged in Titan Krios and tilt series data were collected using software Tomography 5 with dose symmetric strategy with a 3-degree increment from 0 to +/-18 degree. Each tilt was collected as a movie stack of three frames with 1 electron per frame at 1.08 Å/pix. The data was collected at a defocus range of -1.5 to -2.5 μm.

### Data processing

The movie stack of each tilt was motion corrected and averaged using Warp^3,29^. And the CTF of each tilt image was estimated with CTFFIND 4^30^. The tilt series were aligned using Aretomo^31^, and tomograms were reconstructed by RELION 4^31,32^. The reconstructed tomograms are then used for particle picking with the EMAN 2 deep learning algorithm^33^. The rest of the data process was performed in RELION 4. In brief, the 4x binned pseudo-subtomograms were extracted and initial models were built as the reference for 3D classification to sort out the false positive particles and particles as octadecameric complexes. The hexadecameric complex particles were aligned at binning factor 4 and 2 before being refined with no binning. Tomo CTF refinement and frame alignment were performed after the auto-refinement, achieving the final resolution of 2.56 Å with D8 symmetry imposed.

### Cumulative dose analysis and ice thickness analysis

Maps were reconstructed in RELION 4 from the same set of hexadecameric particles but of only one tilt without further alignment. Particles per tomogram were plotted to estimate the ice shape and thickness. Ice thickness per particle was defined by the electron path length and estimated to be 200 Å (the rough MmCpn diameter) plus the Z difference between the top and bottom particle in each of 5 by 5 bins on the XY plane of the tomogram. Particles were grouped based on the ice thickness and the map was reconstructed per ice thickness level with reference being low pass filtered to 15 Å. For each ice thickness, the particle sub-dataset was randomized before subsampling triple to six times without duplicates among the subsampled datasets or within each sub-dataset. And maps were reconstructed for each sample size per ice thickness.

### Model building and refinement

An MmCpn subunit model in the closed form (not published) was fit into the 2.56 Å map and optimized with multiple software including ISOLDE^33,34^,COOT^35^, and deep learning algorithm (Muyuan Chen, manuscript in preparation). The full model was then built from the subunit model by symmetry in Phenix^36^. Figures are prepared using Chimera^37^ and ChimeraX^38^.

## Acknowledgements

We acknowledge Muyuan Chen’s help with model refinement. We acknowledge funding support from Silicon Valley Community Foundation CZI Imaging Initiative, and the Stanford School of Medicine Dean’s Postdoctoral Fellowship.

## Author contributions

Y.Z. and W.C. conceived the project and approach. Y.Z. planned the experiments, collected and analyzed the data. Y.Z. took the lead in writing the manuscript. M.S., W.C. provided critical feedback and helped shape the research, analysis and manuscript.

## Data accessibility

The cryo-EM maps and model will be accessible in EMDB and PDB (accession numbers:EMDBxxx shown in Figure 1, S1, and 2 and PDB yyyy shown in Figure 1 respectively).

## Declaration of interests

The authors declare no competing interests.

---

**Figure S1.**
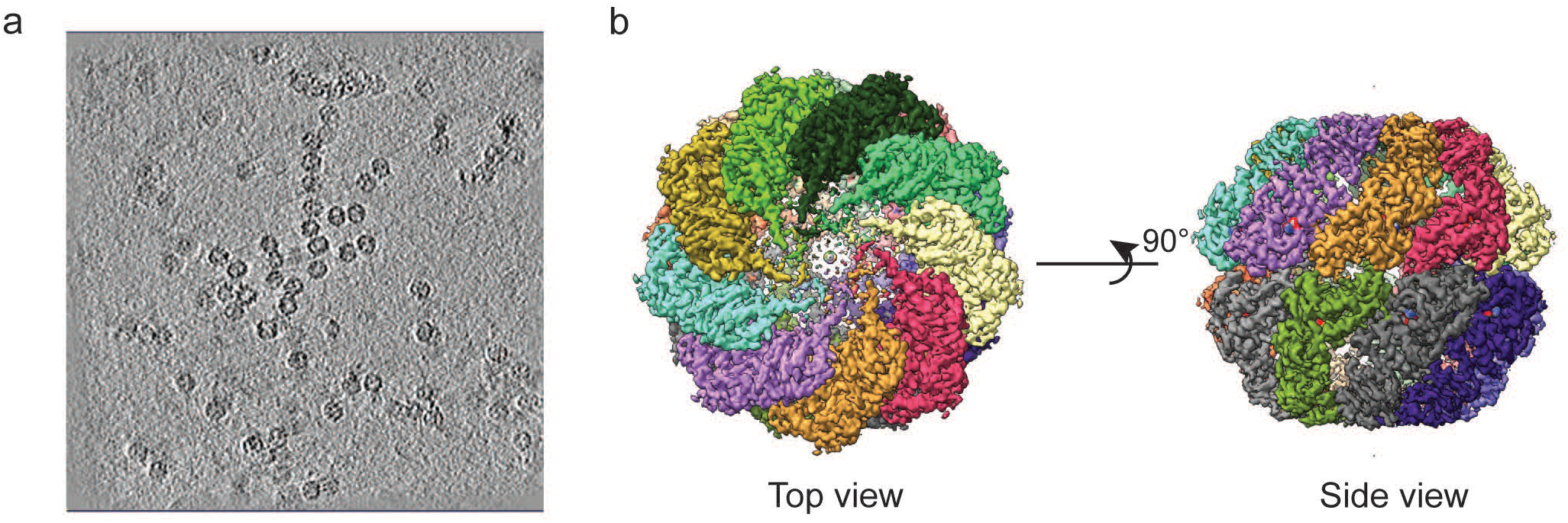
(a) An example slice of a tomogram with the MmCpn molecules embedded in the ice. (b) Reconstruction of a subset of octadecameric MmCpn images in the closed conformation with D9 symmetry.

**Figure S2.**
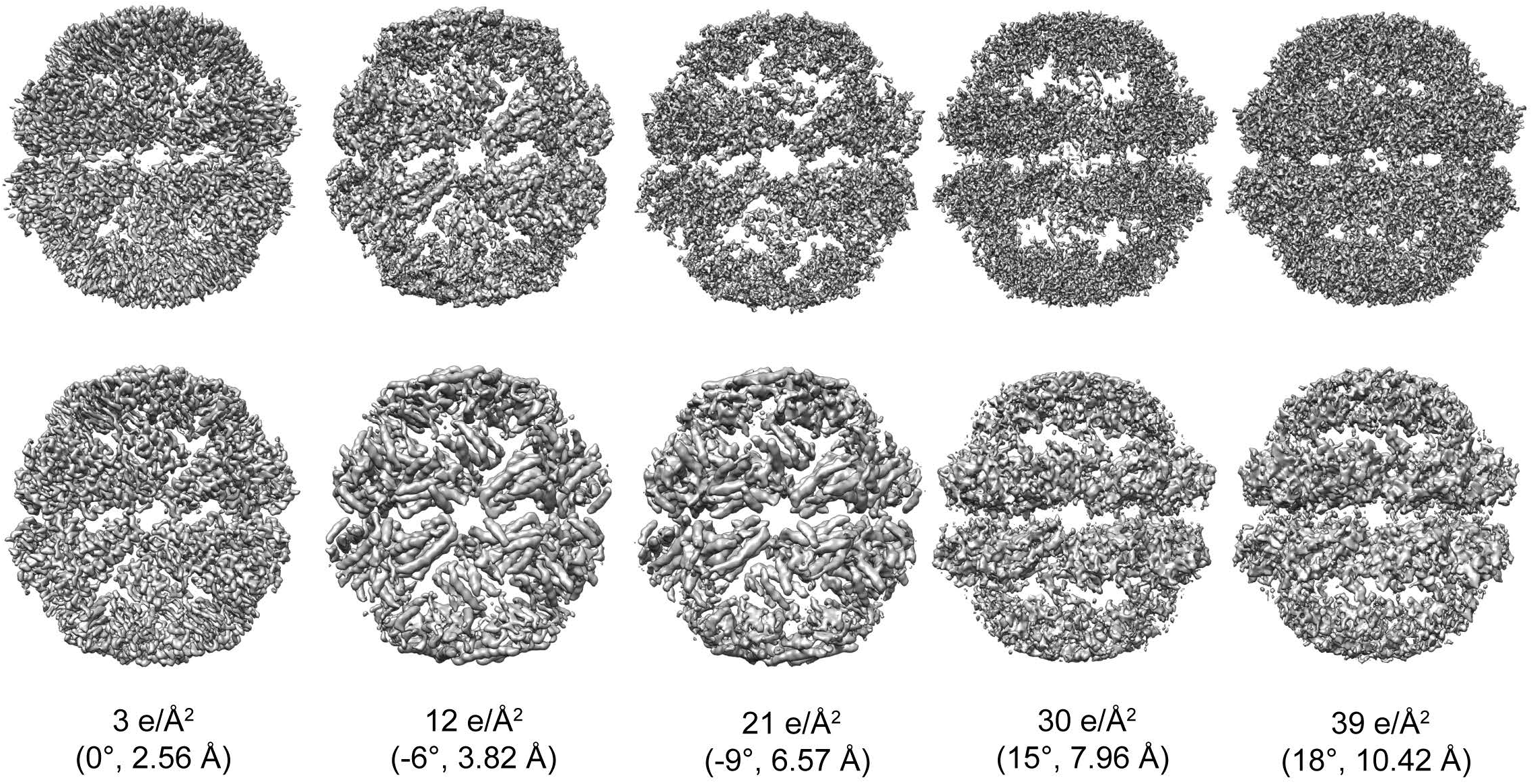
Top row shows the full map of MmCpn reconstructed from the 158,666 subtomograms at a specific tilt angle. Bottom row shows the gaussian filtered maps from the top row.

**Figure S3.**
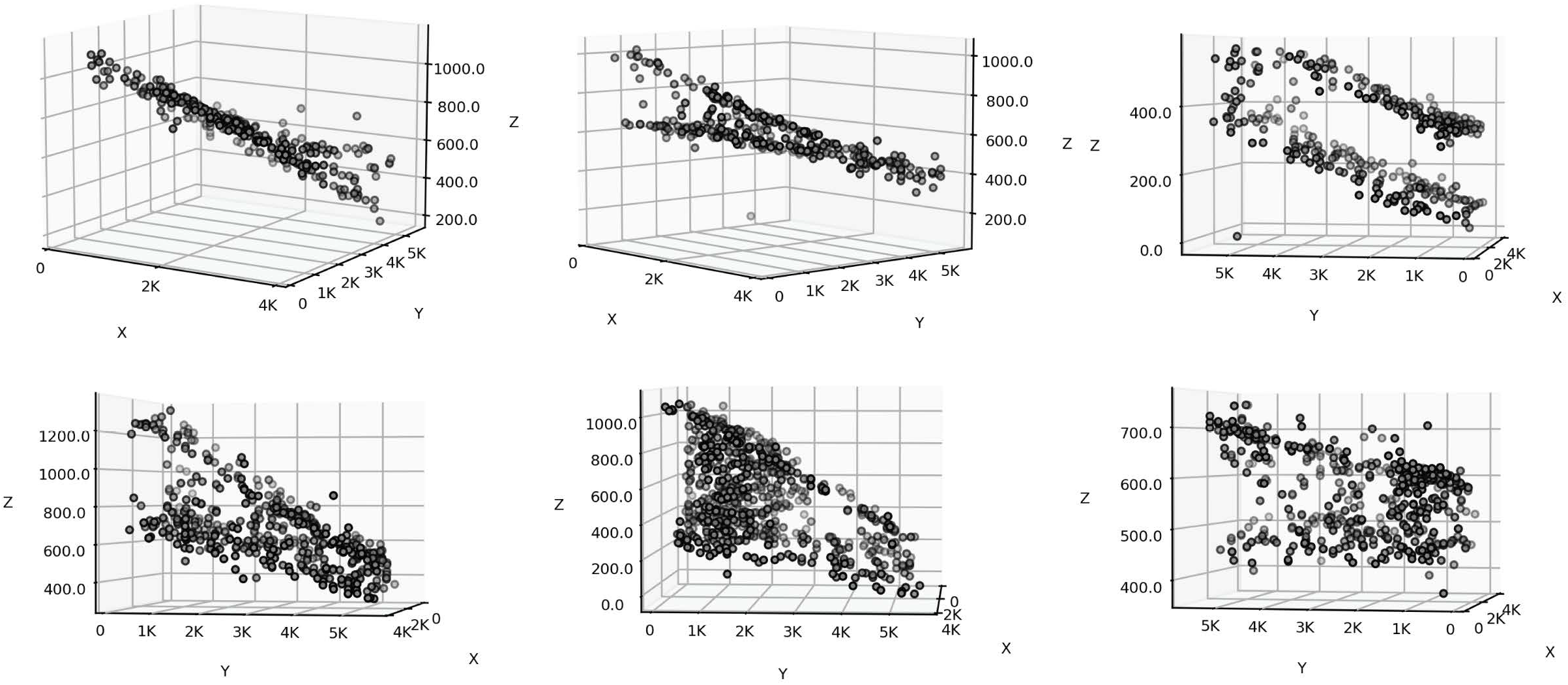
The examples of macromolecule distribution in vitreous ice of various thickness. The gray spots are MmCpn molecules embedded in the vitreous ice.

## Notes

### Competing Interest Statement

The authors have declared no competing interest.

### Summary of Updates

Reformat

